# Mass spectrometry-based proteomic and glycoproteomic drafts of fifteen human body fluids

**DOI:** 10.1101/2025.09.29.679374

**Authors:** Fei Cai, Yingying Ling, Qingyuan Zheng, Ling Li, Tao Su, Jing Shen, Zihan Wu, Meng Gong, Hao Yang, Wanjun Zhao, Yong Zhang

## Abstract

The study of proteins and *N*-glycosylation in human body fluids (HBFs) is crucial for understanding both physiological and pathological states, offering valuable insights for disease diagnostics and monitoring. In this study, we present a comprehensive proteomics and glycoproteomics dataset featuring 15 types of precious HBFs (plasma (P), urine (U), cerebrospinal fluid (CSF), pleural fluid (PF), ascitic fluid (AF), synovial fluid (SF), pericardial fluid (PCF), peritoneal dialysis effluent (PDE), bile (B), gastric juice (GJ), seminal plasma (SP), saliva (SA), pancreatic juice (PJ), sweat (SW), and tear (T)). We also provide detailed protocols for protein processing specific to each type of HBF. The resulting peptides and enriched intact *N*-glycopeptides were analyzed via liquid chromatography-tandem mass spectrometry (LC-MS/MS) in data independent acquisition (DIA) and combined electron transfer/higher-energy collisional dissociation and stepped collision energy/higher-energy collisional dissociation (EThcD-sceHCD) mode, respectively. The resulting spectra and searched files have been uploaded to the online repository iProX (www.iprox.cn) and made easily accessible (PXD068799) to facilitate further research into HBF proteomics and glycoproteomics.

## Background & Summary

Substantial advances have been achieved in recent years in the development of mass spectrometry (MS) instruments and proteomics techniques(*1*). MS-based proteomic approaches have emerged as powerful tools for investigating biological systems, offering critical insights into protein abundance profiles and the landscape of post-translational modifications (PTMs). In particular, glycosylation, a common PTM, plays crucial roles in protein folding, stability, and function, influencing various biological processes and disease mechanisms(*2*). Notably, high-resolution MS has facilitated the identification and characterization of more than ten thousand proteins derived from diverse human tissues, yielding an extensive human proteome catalog(*3, 4*). These resources are poised to complement existing human genome and transcriptome datasets, thereby accelerating biomedical research focused on understanding health and disease mechanisms(*5*). Despite this progress, comprehensive proteomic and glycoproteomic maps covering the full spectrum of biofluids or human body fluids (HBFs) are currently lacking.

Clinically, dozens of HBFs are utilized for disease diagnosis, with plasma (P) and urine (U) serving as well-established samples for routine clinical testing. The protein compositions and associated PTMs of these two fluids have been extensively investigated(*6, 7*). However, comprehensive analyses of protein and glycosylation in other HBFs (cerebrospinal fluid (CSF), pleural fluid (PF), ascitic fluid (AF), synovial fluid (SF), pericardial fluid (PCF), peritoneal dialysis effluent (PDE), bile (B), gastric juice (GJ), seminal plasma (SP), saliva (SA), pancreatic juice (PJ), sweat (SW), and tear (T)) remain scarce in the current literature(*8*). Despite having previously conducted proteomic and glycoproteomic studies on a few HBFs (such as P, U, and SP), we have yet to systematically process and characterize all 15 types of HBFs via consistent sample processing and mass spectrometry analysis methods for proteomic and glycoproteomic studies(*9-11*).

To bridge these gaps and establish a basic proteomic and glycoproteomic profile across different types of HBFs, we collected 15 types of precious HBF samples from clinical practice (Fig. 1a). For each pooled HBF from four individuals, we included a comprehensive description of the preprocessing protocols and the use of a consistent, modified filter-aided sample preparation (FASP) method. The peptides obtained were analyzed via data-independent acquisition mass spectrometry (DIA-MS), whereas the intact *N*-glycopeptides (IGPs) enriched via zwitterionic hydrophilic interaction liquid chromatography (ZIC-HILIC) were subjected to electron-transfer/higher-energy collisional dissociation (EThcD) and stepped collision energy/higher-energy collisional dissociation (sceHCD) mass spectrometry (EThcD-sceHCD-MS/MS) (Fig. 1b)(*12, 13*). This analysis process was repeated three times. Finally, we utilized professional software and bioinformatics approaches for data analysis and presentation (Fig. 1c). These methods presented in this study serve as a reference for examining the variations in HBF protein and glycosylation across different diseases. The extensive nature of these proteomic and glycoproteomic datasets offers numerous opportunities for analysis through different methods, ultimately providing new insights into the evolving trends of protein and glycosylation in HBFs.

**Fig. 1.**
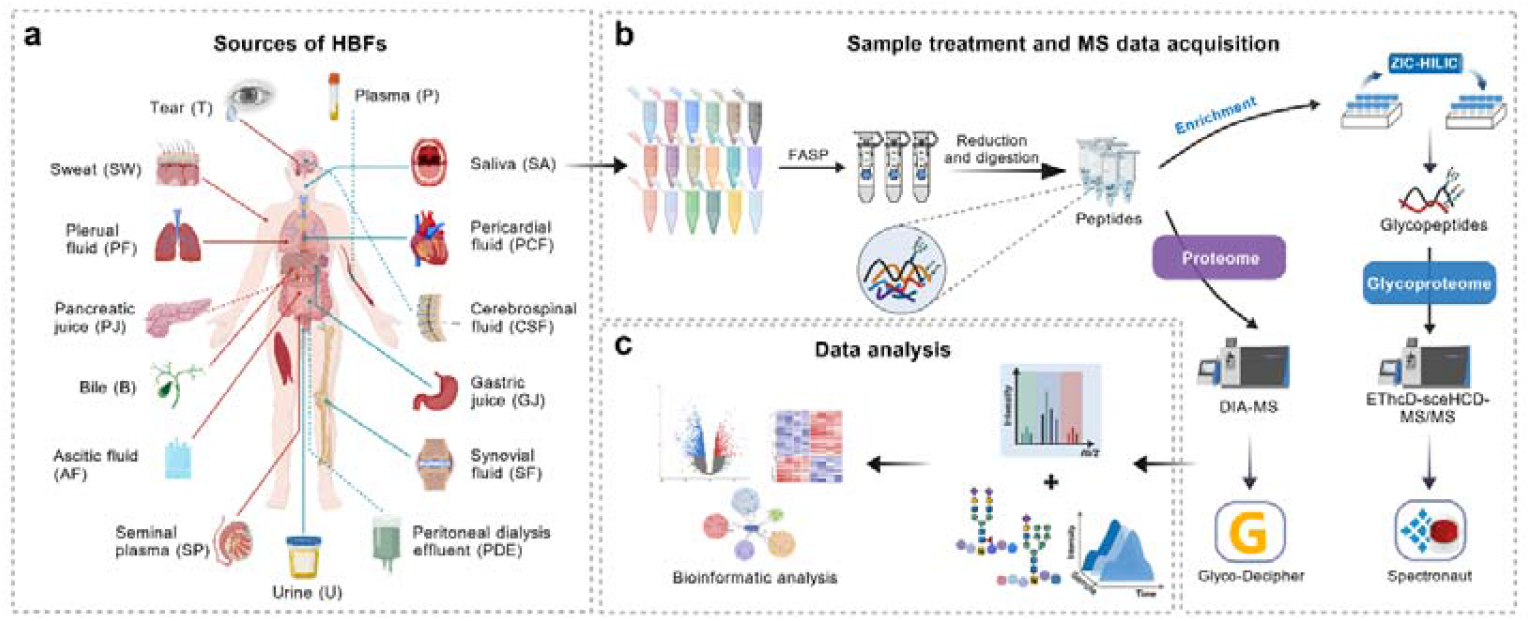
Schematic overview of the fifteen types of HBF samples, along with the corresponding sample treatment and data analysis workflow. **(a)** This panel outlines the sources of the fifteen HBFs investigated in this study. **(b)** The FASP method was employed for all the HBFs. The obtained peptides were analyzed via DIA-MS, whereas IGPs enriched by ZIC-HILIC were subjected to EThcD-sceHCD-MS/MS. **(c)** Bioinformatics pipelines were applied to facilitate data analysis and visualization of the results. Figure created with BioGDP (https://BioGDP.com).

## Methods

### Ethics statement

Pooled human body fluid (HBF) samples (plasma (P), urine (U), cerebrospinal fluid (CSF), pleural fluid (PF), ascitic fluid (AF), synovial fluid (SF), pericardial fluid (PCF), peritoneal dialysis effluent (PDE), bile (B), gastric juice (GJ), seminal plasma (SP), saliva (SA), pancreatic juice (PJ), sweat (SW), and tear (T)) used in this study were selected from several healthy individuals and patients undergoing examination at the Department of Laboratory Medicine, West China Hospital, Sichuan University (Chengdu, China). This study adhered to the principles of the Declaration of Helsinki and received approval from the Ethics Committee of West China Hospital, Sichuan University (Approval No. 2020(102)). Written informed consent was obtained from all participants involved in this study.

### HBF collection and preprocessing

Different HBF samples require different collection and preprocessing methods. **Plasma:** We collected blood samples in EDTA tubes, which were gently inverted four to five times before being centrifuged at 3,000×g and 4□°C for 10 minutes. The P samples were collected and frozen at −80□°C. **Urine:** We gathered random clean-catch midstream U samples in sterile containers and immediately placed them on ice. The samples were then centrifuged at 2,000×g for 30 minutes at 4□°C to remove sediments such as cells and crystals. The supernatant was transferred to clean microcentrifuge tubes, and the resulting urine was subsequently concentrated via ultrafiltration centrifugal devices with a 30-kDa molecular weight cut-off (MWCO). The concentrated urine aliquots were stored at −80□°C for further analysis. **Cerebrospinal fluid:** CSF samples were obtained under aseptic conditions via lumbar puncture. After collection, the samples were immediately centrifuged at 2,000×g for 10 minutes at 4□°C to remove cells and impurities. The clarified fluid was then divided into aliquots in microcentrifuge tubes and stored at −80 °C until analysis. Pleural fluid. PF samples were collected as part of routine clinical thoracentesis or drainage procedures, which were conducted aseptically by experienced physicians via standard techniques. After an adequate volume was collected for clinically necessary tests, the remaining fluid was aseptically transferred into collection tubes. The samples were then centrifuged at 2,000×g for 5 minutes at 4 °C to remove cells and debris. The cell-free supernatant was aliquoted into microcentrifuge tubes and stored at −80 °C for later analysis. **Ascitic fluid:** AF samples were collected during standard clinical procedures, such as abdominal paracentesis or drainage. After collection, the samples were centrifuged at 1,500×g for 5 minutes at 4 °C to eliminate cells and debris. The clear supernatant was then divided into microcentrifuge tubes and stored at −80 °C. **Synovial fluid:** SF samples were aseptically collected from patients through clinical arthrocentesis. After collection, hyaluronidase (e.g., 100 U/mL) was used to reduce the viscosity of the sample, which was incubated it at 37□°C for 30 minutes. The mixture was then centrifuged at 10,000×g for 10 minutes at 4□°C to remove cells and debris. The resulting supernatant was divided into aliquots and stored at −80□°C. **Pericardial fluid:** PCF samples were collected through ultrasound-guided pericardiocentesis. After collection, the samples were centrifuged at 1,500×g for 5 minutes at 4□°C to remove cells and debris. The clear supernatant was then divided into microcentrifuge tubes and stored at −80□°C for subsequent analysis. **Peritoneal dialysis effluent:** PDE samples were collected via catheter drainage during clinical peritoneal dialysis. Following collection, the samples were centrifuged at 1,500×g for 5 minutes at 4□°C to eliminate cells and debris. The cell-free supernatant was then divided into microcentrifuge tubes and stored at −80□°C. **Bile:** B samples were acquired either during clinical procedures or through catheter drainage. Once collected, the samples were mixed with an equal volume of phosphate buffer solution (PBS) to decrease their viscosity. The samples were then centrifuged at 13,000×g for 10 minutes at 4□°C to remove cells and debris. The supernatant was aliquoted into microcentrifuge tubes and stored at −80□°C for future analysis. **Gastric juice:** GJ samples were collected through a gastric tube during gastroscopy. After collection, the samples underwent initial centrifugation at 1,500×g for 5 minutes at 4□°C to remove solid particles. The supernatant was then divided into microcentrifuge tubes and stored at −80□°C. **Seminal plasma:** Semen samples were obtained through masturbation following a period of 3–7 days of sexual abstinence. The samples were then left to fully liquefy for 20 minutes at 37□°C. After liquefaction, the semen was centrifuged at 1000×g for 20 minutes at 4□°C. The supernatant, which consisted solely of SP, was carefully transferred into sterile microcentrifuge tubes and stored at −80□°C. **Saliva:** Nonsmoking, nonpregnant, nonlactating, and nondiabetic volunteers were screened for good general and oral health with normal salivary function. Importantly, none of them were taking any medications. Unstimulated SA samples were collected between 9 a.m. and 10 a.m., at least two hours after the last intake of food. Before collection, the participants rinsed their mouths with physiological saline. The SA samples were then collected and placed on ice. A protease cocktail inhibitor was added to the samples immediately after collection to minimize protein degradation. After centrifugation at 12,000×g at 4 °C for 10 minutes to remove insoluble materials, the resulting supernatant was collected and stored at −80 °C. **Pancreatic juice:** PJ samples were collected either during endoscopic retrograde cholangiopancreatography (ERCP) or through surgical exposure of the pancreatic duct. Each sample was immediately placed on ice and treated with protease inhibitors. To remove mucus, cellular debris, and other particulates, the PJ samples were centrifuged at 12,000×g for 15 minutes at 4□°C. The clarified supernatant was transferred into clean microcentrifuge tubes and stored at −80□°C. **Sweat:** Volunteers were instructed to avoid using cosmetics and contact chemical substances for 24 hours prior to sampling. SW samples were collected during exercise (running) via sterile gauze pads pressed against the skin. These gauze pads were stored in precooled sterile 50-mL centrifuge tubes and placed on ice. After 5 to 6 gauze pads were collected from each participant, the samples were taken to the laboratory. The SW samples were extracted by centrifuging the gauze at 12,000×g for 15 minutes at 4□°C, which resulted in an average volume of approximately 4 mL per participant.

To remove cells, keratins, and other impurities, the SW samples were further centrifuged at 13,000×g for 10 minutes at 4□°C. The supernatant was then concentrated via 30-kDa ultrafiltration devices and stored at −80□°C. **Tear**. A polyester wick was used to collect T samples by placing them at the tear meniscus along the lower eyelid margin. The samples were then transferred to microcentrifuge tubes, and T samples were extracted from the polyester wick by centrifuging each sample at 3,000×g for 10 minutes at 4□°C. The supernatant was stored at −80□°C.

### SDS-PAGE analysis

Sodium dodecyl sulfate polyacrylamide gel electrophoresis (SDS□PAGE) analysis was conducted via the following procedure: These samples were boiled for 10 minutes following the addition of a 5×loading buffer. After centrifuging at 13,000×g for 15 minutes, these proteins were loaded onto 12% SDS□PAGE gels. Electrophoresis commenced at 100 V at 4 □. After 20 min, the voltage was increased to 120 V, and electrophoresis was continued for an additional hour. Once the dye front exited the gel, the lanes were excised and stained with the BluPower Fast Staining Coomassie. Water was then used for decolorization.

### Protein lysis and digestion

The proteins (200 μg) from 15 HBF samples subjected to proteolysis were subjected to a modified filter-aided sample preparation (FASP) protocol. Initially, the proteins were placed into a 30-kDa filter and then centrifuged at 25 °C at 13,000×g for 15 min. Subsequently, 200 μL of 20 mM dithiothreitol (DTT) was added to the mixture, which was allowed to react for 4 hours at 37 °C. Next, 200 μL of 50 mM iodoacetamide (IAA) was added, and the resulting mixture was incubated in the dark at room temperature for 1 hour. The urea (UA) buffer was then replaced with 50 mM ammonium bicarbonate (NH□HCO□) buffer by centrifuging three times at 13,000×g for 15 minutes. Afterward, 4 μg of trypsin was added, and digestion was performed overnight at 37 °C. Once digestion was complete, the peptides were collected by centrifugation at 13,000 g for 15 minutes. The peptide concentration was determined via a quantitative colorimetric peptide assay. Finally, the peptide samples were freeze-dried via a SpeedVac system (Thermo Fisher Scientific, USA).

### Intact *N*-glycopeptide enrichment

The lyophilized peptides (100 μg) were resuspended in 70 μL of binding buffer (80% ACN/0.2% TFA). The intact *N*-glycopeptides were subsequently enriched via zwitterionic hydrophilic interaction chromatography (ZIC-HILIC) following previously described procedure(*14*). Specifically, 100 μg of tryptic peptides were mixed with 5 mg of ZIC-HILIC materials in binding buffer (80% ACN/0.2% TFA). After incubation at room temperature with rotation for 2 hours, the mixture was transferred to a pipet tip containing a C8 membrane. The hydrophobic peptides were then washed with binding buffer, and the IGPs were eluted with 70 μL of 0.1% TFA solution. The eluent was dried by SpeedVac for further analysis.

### LC-MS/MS analysis

Peptide and IGP analyses were performed via an Orbitrap Fusion Lumos mass spectrometer (Thermo Fisher, USA), which was connected to an EASY-nL 1200 system and a nanospray ion source. The liquid chromatography (LC) gradient employed buffer A (0.1% FA) and buffer B (80% ACN with 0.1% FA). The samples were resuspended in 10 μL of buffer A, and 1 μL of this mixture was injected into and separated on a column (ReproSil-Pur C18-AQ, with a particle size of 1.9 μm, an inner diameter of 100 μm, and a length of 25 cm; supplied by Dr Maisch) over a 78-min gradient (0−8 min, 5−12% B; 8−58 min, 12−22% B; 58−70 min, 22−32% B; 70−71 min, 32−90% B; and 71−78 min, 90% B) at a flow rate of 350 nL/min. The column was maintained at 55 °C. The parameters for the proteomics analysis were configured as follows: The MS operated in direct data-independent acquisition (DIA) mode. Each cycle included one full MS scan along with 60 DIA scan windows, covering a mass range from 350 to 1,500 m/z. The operational parameters were finely tuned: the full MS scan resolution was set to 60,000 at 200 m/z; the automatic gain control (AGC) target was established at 1e6; the maximum injection time (MIT) was capped at 50 ms; and the scans were executed in profile mode. The DIA scans were conducted at a resolution of 15,000, with an AGC target of 5e5, an MIT of 22 ms, and high-energy collision dissociation (HCD) mode with a normalized collision energy (CE) of 30%, operating in centroid mode. The parameters for glycoproteomic analysis were as follows: detection was carried out in the data-dependent acquisition (DDA) mode. The detailed parameters have been described previously(*10, 15, 16*). Specifically, this analysis alternated between EThcD and sceHCD modes in a 3-second cycle. During the 2-second EThcD phase, MS1 spectra were gathered in the 400–1600 m/z range at a resolution of 60,000. The settings for the RF lens, AGC target, MIT, and exclusion duration were adjusted to 40%, 2e5, 50 ms, and 15 s, respectively. MS2 spectra were captured with a 2 m/z isolation width and a resolution of 30,000. The AGC target, MIT, and EThcD type settings were 5e5, 150 ms, and 35%, respectively. During the 1-second sceHCD phase, MS1 spectra were again recorded across a 400–1600 m/z range at a 60,000 resolution. The RF lens, AGC target, MIT, and exclusion duration were set to 40%, standard, auto, and 15 s, respectively. MS2 spectra of precursor ions were selected with a 2 m/z isolation width and obtained at a resolution of 30,000, with the AGC target set to 200%, the MIT to 150 ms, and the stepped HCD collision energy set to 20-30-40%.

### Data search

Proteomics data files were searched against the human UniProt database (version 2015_03, 20,410 entries) using the Spectronaut (v15.4) software which was operated with the default settings unless otherwise specified. The glycoproteomics data files were searched against the same database via Glyco-Decipher (v1.0.5) software(*17, 18*). The mass tolerance parameters were set at ±6 ppm for precursors and ±20 ppm for fragment ions, allowing up to two missed cleavages. The fixed modification in place was carbamidomethyl (C), while oxidation (M) and acetylation (protein N-terminus) were considered variable modifications. The glycan database used was the default.

## Data Records

Both the raw mass spectrometry data (.raw folders generated by Orbitrap Fusion Lumos) and the database search results from the proteomics data (.txt and.tsv files generated by Spectronaut) and glycoproteomics data (.txt files generated by Glyco-Decipher) have been uploaded to iProX as a part of the public dataset PXD068799.

## Technical Validation

### Proteomic profiling of HBFs

To visually assess the differences in protein expression among various HBFs, we performed SDS-PAGE analysis. The results revealed variations in protein expression across the different HBFs, including highly abundant immunoglobulins (Fig. 2a). Using DIA-MS technology, we quantified 4,389 proteins from 15 HBFs in three repeated tests. Furthermore, we can clarify the types and abundances of proteins present in different HBFs. The variety of proteins detected in these HBFs differs (Fig. 2b). For example, on average, 3,275 proteins can be identified in U samples, and 3,135 can be identified in SP samples. In contrast, only 489 proteins were detectable in the P samples. This disparity is primarily because the dynamic range of plasma protein concentrations spans a dynamic range of at least 12 orders of magnitude, and a small subset of high-abundance proteins (HAPs) constitute more than 90% of the total protein mass in plasma(*19, 20*). Furthermore, we illustrate the distribution of unique proteins among these HBFs (Fig. 2c). There are 223 shared proteins that are expressed across all 15 types of HBFs, despite variations in their expression levels. Additionally, we identified proteins uniquely expressed in specific HBFs, such as ANLN in PF, ABCA8 in AF, and A20A4 in PCF. The absence of specifically expressed proteins in plasma might be attributed to its circulatory nature throughout the body. The presence of HBF-specific expressed proteins could be due to limited sample sizes, variation in protein expression abundance, and inherent limitations in MS-based detection capabilities. Therefore, in the field of HBF proteomics, it is crucial to utilize a large number of clinical samples and apply a consistent mass spectrometry method for robust comparative analyses.

**Fig. 2.**
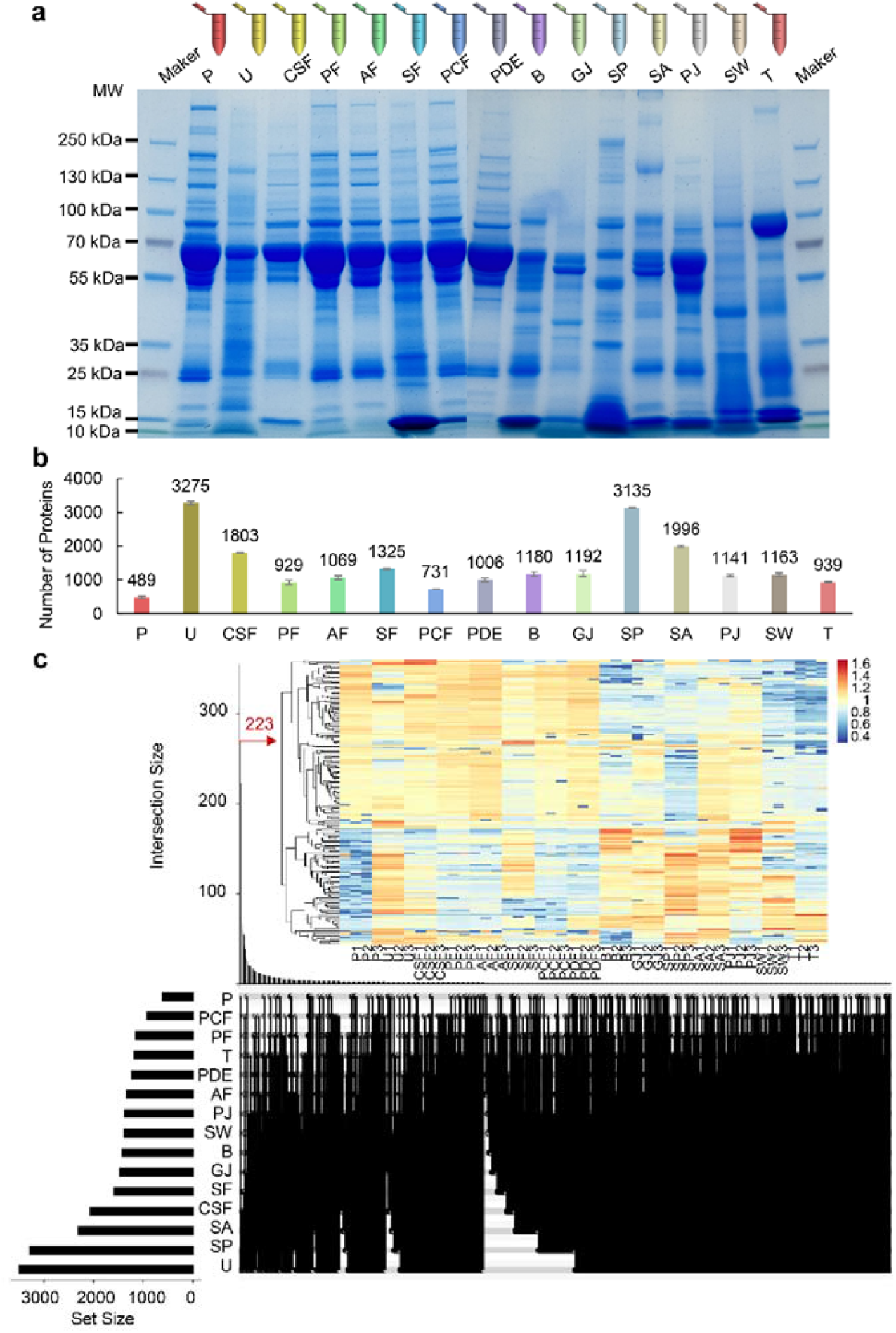
Proteomic characteristics of the fifteen types of HBF samples. **(a)** SDS-PAGE analysis displays variations in protein expression across different HBF samples. **(b)** Bar chart presenting the quantified protein counts from these 15 HBF samples as determined by DIA-MS. **(c)** UpSet plot showing the distribution of unique proteins among these samples. Furthermore, heatmap analysis of the 223 shared proteins across the 15 HBF samples revealed significant differences in their expression levels.

### Glycoproteomic profiling of HBFs

Across the 15 HBF samples analyzed, a total of 7,827 unique IGPs were quantified, corresponding to 617 distinct *N*-glycopeptide sequences, 730 unique *N*-glycan compositions, and 349 *N*-glycoproteins. Compared with the proteomic dataset, the inter-HBF differences were further exacerbated at the IGP level. Similarly, the qualitative and quantitative reproducibility of three replicates was lower than that of the proteomic data, which can be attributed to the complexity of sample processing workflows and the inherent challenges of IGP analysis (Fig. 3a). Nevertheless, the number of quantified IGPs exhibited a trend largely consistent with that of the proteomic data. Additionally, heatmap analysis of the 36 IGPs with high quantitative detection rates (>50%) across the 15 HBF samples revealed significant differences in their expression patterns.

**Fig. 3.**
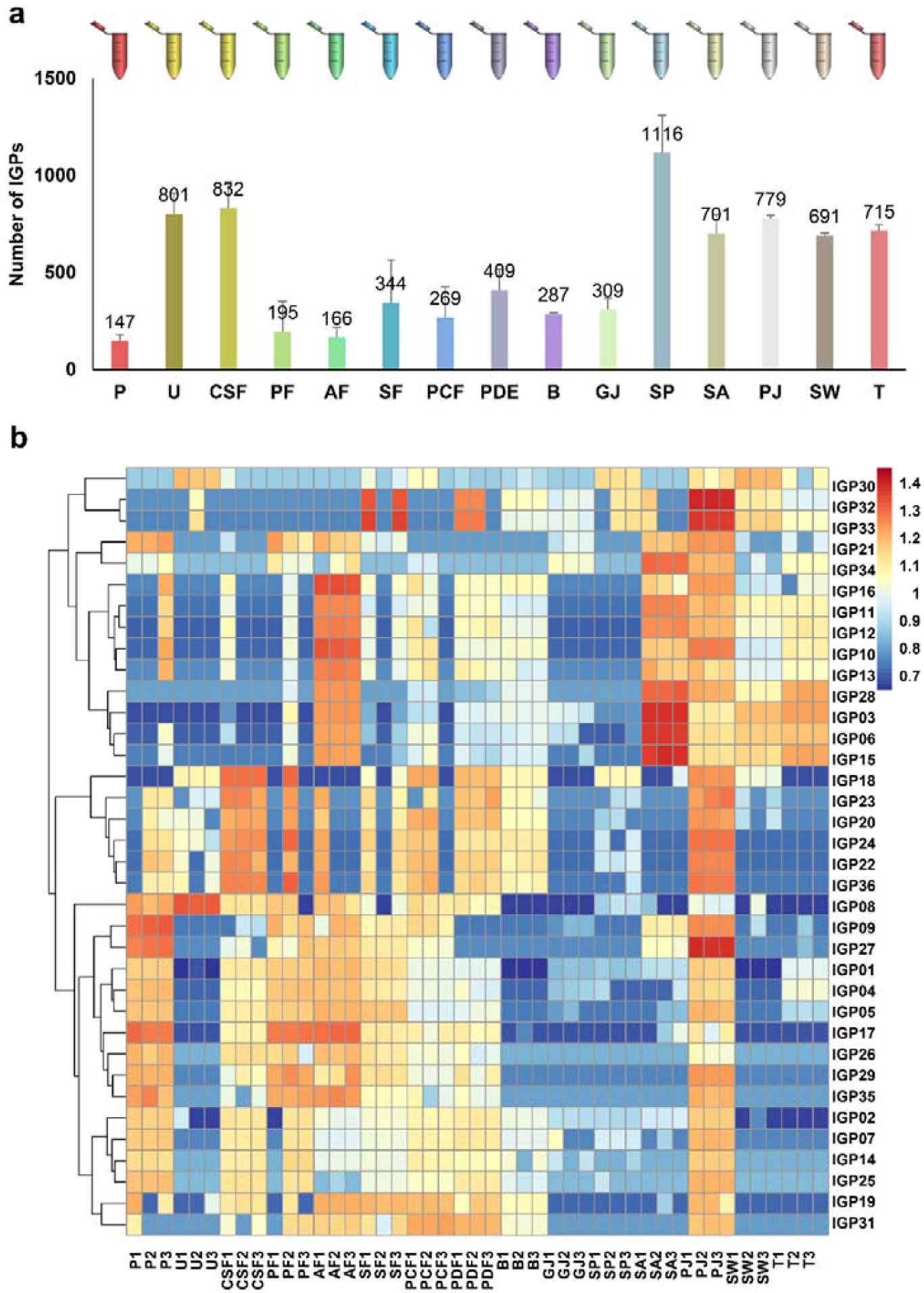
Glycoproteomic characteristics of the fifteen types of HBF samples. **(a)** Bar chart displaying the quantified IGP counts from these 15 HBF samples as determined by EThcD-sceHCD-MS/MS. **(b)** Heatmap analysis of the 36 IGPs with high quantitative detection rates (>50%) across the 15 HBF samples highlights significant differences in their expression levels.

## Usage Notes

The online proteomics and glycoproteomics data include both raw output files and results processed via specific software: Spectronaut (v15.4) for proteomics and Glyco-Decipher (v1.0.5) for glycoproteomics. Researchers have the option to analyze these raw files with different software or settings than those used in this study. Alternatively, they can directly access the processed data. There are various methods for interpreting the information in the resulting output files. Given that mass spectrometry intensities follow a lognormal distribution, applying a logarithmic transformation to these intensities is commonly performed prior to comparative analyses, assuming a normal distribution. The dataset also contains missing values, which can be reduced via various methods.

## Code availability

No custom computer codes were generated in this work. Bioinformatic analysis was performed via the OmicStudio tools at https://www.omicstudio.cn.

## Data availability

The dataset has been deposited to iProX (www.iprox.cn) and is easily accessible (PXD068799).

## Acknowledgements

We are grateful to the volunteers for providing body fluid samples. We thank the Mass Spectrometry Platform of West China Hospital for performing the LC-MS/MS analysis. We are grateful for the editing service provided by the Library of West China Hospital.

## Author contributions

F.C., W.Z. and Y.Z. wrote the manuscript. F.C., Y.L., Q.Z., L.L., J.S. and Z.W. provided the clinical samples and prepared the samples. T.S. performed the mass spectrometry analysis.

Y.Z. performed the data analysis. M.G., H.Y., W.Z. and Y.Z. supervised the project. All the authors contributed to editing the manuscript.

## Funding

The authors greatly acknowledge the financial support from the National Natural Science Foundation of China (Grant No. 92478101) and the National Key R&D Program of China (Grant No. 2022YFF0608401).

## Competing interests

The authors declare no competing interests.

## Additional information

**Correspondence** and requests for materials should be addressed to Y.Z.

**Reprints and permissions information** is available at www.nature.com/reprints.

**Publisher’s note** Springer Nature remains neutral with regard to jurisdictional claims in published maps and institutional affiliations.

## Notes

https://www.iprox.cn//page/SCV017.html?query=PXD068799

### Competing Interest Statement

The authors have declared no competing interest.

## References

1. Y. Zhao et al., Evolution of Mass Spectrometry Instruments and Techniques for Blood Proteomics. Journal of proteome research 22, 1009–1023 (2023).

2. K. Thaçi, R. M. Anthony, The importance of IgG N-glycosylation in Health, Disease, and Neonatal hemochromatosis. Glycoscience & Therapy 1, 100002 (2025).

3. M. Wilhelm et al., Mass-spectrometry-based draft of the human proteome. Nature 509, 582–587 (2014).

4. M. S. Kim et al., A draft map of the human proteome. Nature 509, 575–581 (2014).

5. B. F. Cravatt, G. M. Simon, J. R. Yates, 3rd, The biological impact of mass-spectrometry-based proteomics. Nature 450, 991–1000 (2007).

6. J. Rutledge et al., Comprehensive proteomics of CSF, plasma, and urine identify DDC and other biomarkers of early Parkinson’s disease. Acta neuropathologica 147, 52 (2024).

7. P. Caron et al., A liquid chromatography-mass spectrometry assay for the quantification of nucleotide sugars in human plasma and urine specimens and its clinical application. Journal of chromatography. A 1677, 463296 (2022).

8. Y. T. Chen, W. R. Liao, H. T. Wang, H. W. Chen, S. F. Chen, Targeted protein quantitation in human body fluids by mass spectrometry. Mass spectrometry reviews 42, 2379–2403 (2023).

9. Y. Zhang et al., Characterization of N-linked intact glycopeptide signatures of plasma IgGs from patients with prostate carcinoma and benign prostatic hyperplasia for diagnosis pre-stratification. The Analyst 145, 5353–5362 (2020).

10. T. Lin et al., Characterization of site-specific N-glycosylation signatures of isolated uromodulin from human urine. The Analyst 148, 5041–5049 (2023).

11. G. Yan et al., GlycoIP: an integrated platform for simultaneous and site-specific N/O-glycosylation analysis of human semen. Frontiers in chemistry 13, 1569561 (2025).

12. Y. Mao, Y. Zhao, Y. Zhang, H. Yang, In-depth characterization and comparison of the N-glycosylated proteome of two-dimensional- and three-dimensional-cultured breast cancer cells and xenografted tumors. PloS one 15, e0243789 (2020).

13. Y. Zhao et al., A Novel Integrated Pipeline for Site-Specific Quantification of N-glycosylation. Phenomics (Cham, Switzerland) 4, 213–226 (2024).

14. Y. Zhang et al., Site-specific N-glycosylation Characterization of Recombinant SARS-CoV-2 Spike Proteins. Molecular & cellular proteomics : MCP 20, 100058 (2021).

15. Y. Zhang et al., Sequential Analysis of the N/O-Glycosylation of Heavily Glycosylated HIV-1 gp120 Using EThcD-sceHCD-MS/MS. Frontiers in Immunology 12, 755568 (2021).

16. W. Zeng et al., Comparative N-Glycoproteomics Analysis of Clinical Samples Via Different Mass Spectrometry Dissociation Methods. Frontiers in chemistry 10, 839470 (2022).

17. R. Lou et al., Benchmarking commonly used software suites and analysis workflows for DIA proteomics and phosphoproteomics. Nature communications 14, 94 (2023).

18. Z. Fang et al., Glyco-Decipher enables glycan database-independent peptide matching and in-depth characterization of site-specific N-glycosylation. Nature communications 13, 1900 (2022).

19. Y. T. Deng et al., Atlas of the plasma proteome in health and disease in 53,026 adults. Cell 188, 253–271.e257 (2025).

20. G. H. Eldjarn et al., Large-scale plasma proteomics comparisons through genetics and disease associations. Nature 622, 348–358 (2023).

